# Self-assembled DNA-collagen bioactive scaffolds promote cellular uptake and neuronal differentiation

**DOI:** 10.1101/2024.05.24.595848

**Authors:** Nihal Singh, Ankur Singh, Dhiraj Bhatia

## Abstract

Different modalities of DNA-Collagen complexes have been utilized primarily for gene delivery studies. However, very few studies have investigated the potential of these complexes as bioactive scaffolds. Further, no studies have characterized the DNA-Collagen complex formed from the interaction of self-assembled DNA macrostructure and collagen. Towards this investigation, we report herein the fabrication of novel bioactive scaffolds formed from the interaction of sequence-specific, self-assembled DNA macrostructure and collagen type I. Varying molar ratios of DNA and collagen resulted in highly intertwined fibrous scaffolds with different fibrillar thicknesses. The formed scaffolds were biocompatible and presented as a soft matrix for cellular growth and proliferation. Cells cultured on DNA/Collagen scaffolds promoted enhanced cellular uptake of transferrin, and the potential of DNA/Collagen scaffolds to induce neuronal cell differentiation was further investigated. The DNA/Collagen scaffolds promoted neuronal differentiation of precursor cells with extensive neurite growth in comparison to control groups. These novel, self-assembled DNA/Collagen scaffolds could serve as a platform for the development of various bioactive scaffolds with potential applications in neuroscience, drug delivery, tissue engineering, and in vitro cell culture.

**Graphical Abstract:** 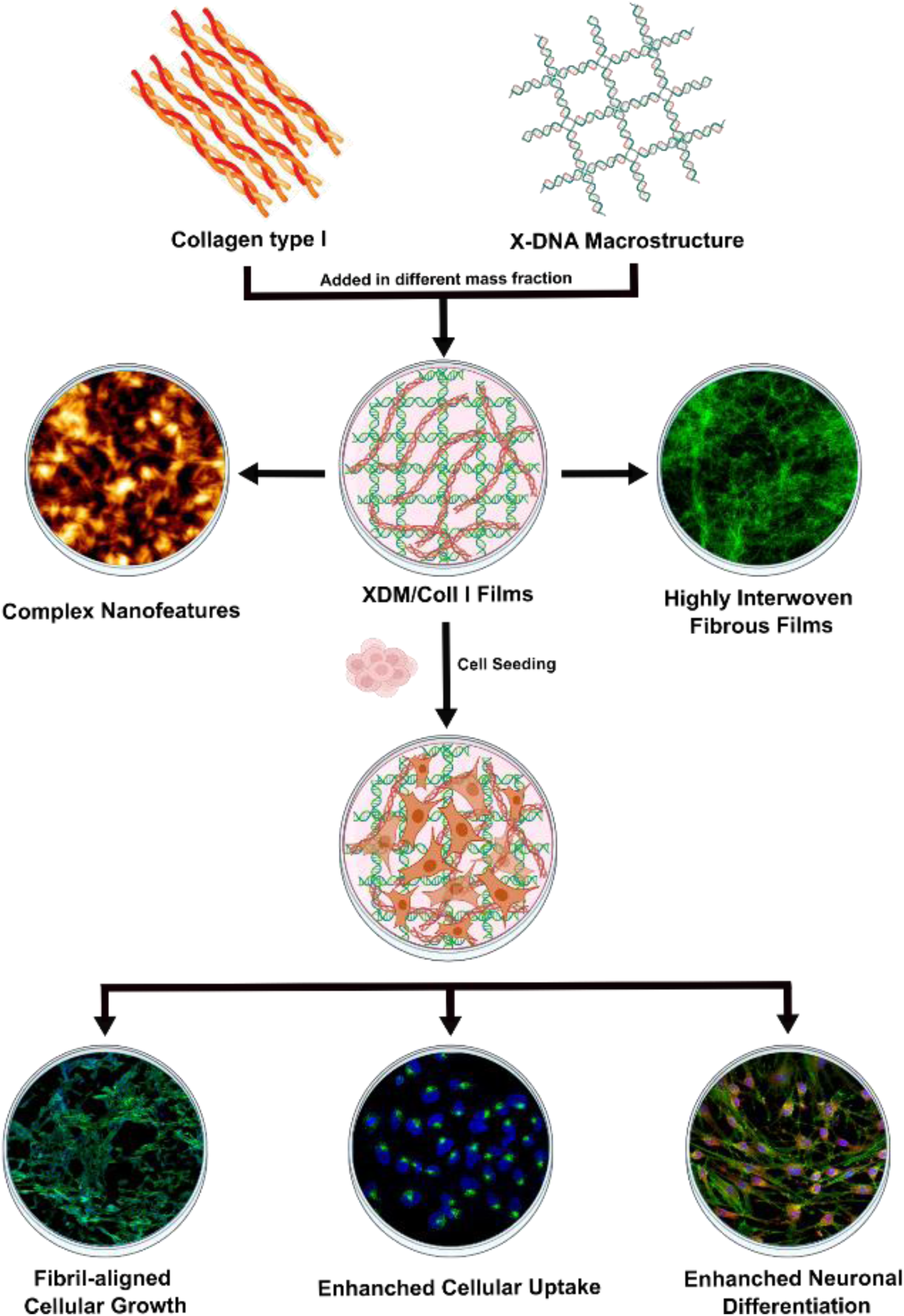

## 1. Introduction

Exploring how DNA and proteins interact has been a major focus of molecular biology research for understanding various cellular processes^1^. With an increased understanding of DNA-protein interactions, these insights were utilized in more translational research fields, including tissue engineering, drug development, and gene editing^2,3^. One such interaction that has been extensively studied, especially from the perspective of gene transfection, is the DNA-collagen interaction^4–6^. First discovered in 1976, researchers identified that both single-stranded DNA (ssDNA) and double-stranded DNA (dsDNA) strongly interacted with collagen and collagen-like materials in the glomerular basement membrane via electrostatic interactions^7^. Due to the nature of interaction involving collagen triple helix surrounding the central core DNA, thereby preventing its degradation, the DNA-collagen complex was extensively utilized as a gene carrier^4,8,9^.

Initial research focused on using DNA-collagen nanoparticle aggregates for their selective uptake and endocytosis in gene transfection investigations^8,10^. However, it was not until 1997 that Kitamura et al. observed the effect of DNA structure on collagen fibrillogenesis^11^. The author observed that the addition of DNA to collagen during fibrillogenesis results in the spontaneous formation of collagen fibrils with a crossbanding pattern. Further investigation by Kaya et al. identified that the fibrillogenesis rate and the resulting fibrils’ structure are influenced by the type of DNA used (linear/cyclic/dsDNA/ssDNA)^12^. Subsequently, the influence of other parameters, including molecular weight, purity, and DNA-collagen mass fraction, on fibril formation was identified^12^. However, most of these studies utilized high molecular weight (>1000 base pairs) random sequence dsDNA, and the effect of small ssDNA (<200 base pairs) on fibril formation was not extensively characterized. In a recent study, James et al. investigated the effect of random sequence ssDNA on fiber formation and observed that the ssDNA length dictates the self-assembly of DNA-collagen fibers, with ssDNA of 15-90 base pair lengths spontaneously forming fibers of various lengths at room temperature^13^.

Despite extensive research characterizing DNA-collagen complexes, there have been no studies investigating how sequence-specific, self-assembled DNA macrostructures affect the formation of DNA-collagen complexes. Moreover, the impact of the relative ratio of DNA macrostructures to collagen on complex formation also remains unexplored. Further, apart from gene delivery applications, very few studies have explored the potential of DNA-collagen complexes for other translative applications. The potential of the DNA-collagen fibrillar complex has been suggested in a few studies, where it has been employed for applications such as wound healing, protein biosensing, and tissue engineering^14–17^. As collagen is widely recognized for its role in regulating various cellular processes like adhesion, proliferation, and differentiation, the potential of the DNA-collagen fibrillar complex as a bioactive coating and scaffolds for in vitro cell culture applications appears extremely promising^18^.

To develop DNA-collagen-based scaffolds for cellular and biomedical applications, herein, we report for the first time the interaction of self-assembled DNA macrostructure with collagen type I to form bioactive scaffolds. We investigated the effect of different mass ratios of DNA and collagen on the bioactive scaffold formation and characterized their structures utilizing Atomic Force Microscopy (AFM) and Scanning Electron Microscopy (SEM). We then investigated the growth and proliferation of breast cancer cell line (SUM159), on the DNA/collagen scaffolds. Further, we also evaluated various cellular processes, including cellular adhesion and endocytic uptake on these scaffolds. Lastly, to probe the differentiation potential of DNA/collagen scaffolds, the differentiation of SH-SY5Y precursor cells into mature neurons was examined.

## 2. Results and Discussions

### 2.1. Synthesis of Self-assembled X-DNA Macrostructure

A branched four-way junction DNA macrostructure, named X-DNA macrostructure (XDM), was utilized to study the interaction between self-assembled DNA macrostructure and collagen. We decided to utilize XDM for our study as it forms an extensive branched DNA network by self-assembly of four ssDNA primers, making it appropriate to study the interaction of collagen with a dense DNA macrostructure network^19^ **(Fig. 1A)**. Electrophoretic mobility shift assay (EMSA) was performed to confirm the assembly of four primers to form XDM. As observed, due to network formation in XDM, its mobility on the gel is reduced compared to single oligo primer (X1), confirming the formation of a higher-order structure **(Fig. 1B)**.

**Fig.1.**
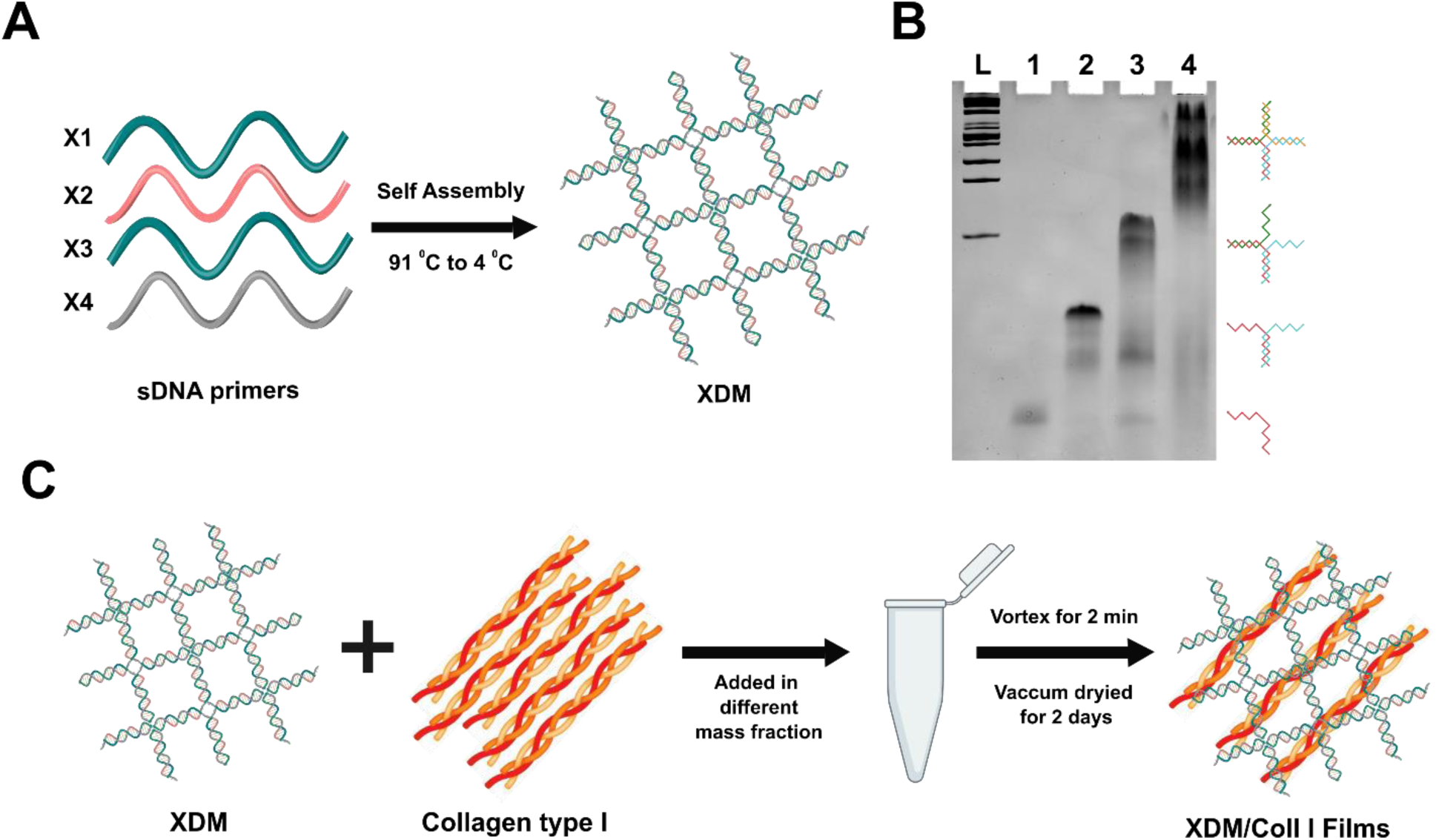
Synthesis of XDM/Coll I scaffolds. (A) Schematic representation of XDM formation from its consequent ssDNA primers (X1, X2, X3, and X4). (B) Electrophoretic mobility shift assay (EMSA) to confirm self-assembly of XDM: L, 100 bp DNA ladder; lane 1, X1; lane 2, X1+X2; lane 3, X1+X2+X3; lane 4, X1+X2+X3+X4 (XDM). (C) Schematic representation for the synthesis of XDM/Coll I scaffolds.

### 2.2. Synthesis of DNA-Collagen Scaffolds

Previous studies have shown that several parameters, including DNA molecular weight, oligomer length, purity, and weight ratio, affect the type and structure of the DNA-collagen complex formed^12^. DNA/Collagen mass fraction, in particular, plays an important role in the fibrillogenesis of the DNA-Collagen complex. Therefore, we investigated the effect of different mass fractions of XDM and collagen type I (XDM / XDM+Coll I;20%, 50%, 90%) on the formation of bioactive scaffolds. The 0% mass fraction was taken as collagen control, and the 100% mass fraction was taken as DNA control. We observed scaffold formation in the cases of 20% and 50% mass fraction; however, no scaffolds were observed in the case of 90% mass fraction. Visualization of scaffolds under the optical microscope showed that the scaffolds were made up of a dense fibrous network, with 20% XDM/Coll I scaffolds containing thinner fibrous networks compared to 50% XDM/Coll I scaffolds **(Fig. 2)**. Suggesting that the mass of XDM increased the thickness of the fibrous network increased. Also, to confirm that the observed fibrous scaffolds were formed because of XDM and Coll I specific interaction, we mixed single oligo primer (X1) at 50% mass fraction with Coll I. Upon mixing, we observed thick individual fibers **(Fig. S1),** which is in line with observation from James et al., where the authors demonstrated that short monodisperse oligonucleotides (<100 base pair) form self-assembled fibers upon complexing with Coll I, suggesting the interaction of XDM with collagen results in extensive fibrous network instead of forming self-assembled individual fibers^13^.

**Fig. 2.**
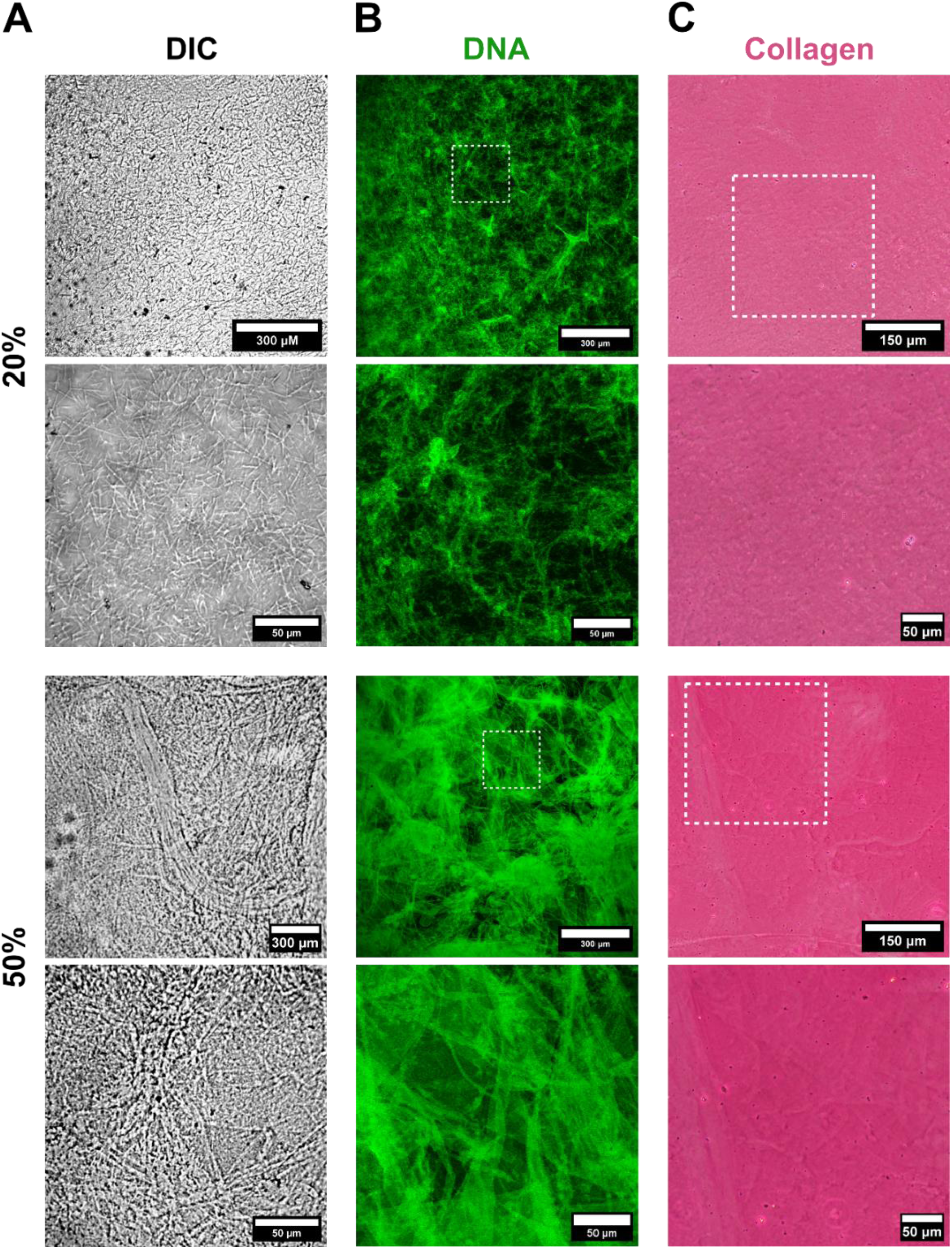
Visualization of the distribution of DNA and collagen in XDM/Coll I scaffolds. The 20% and 50% XDM/Coll I scaffolds were visualized using (A) Differential Interference Contrast (DIC) microscopy at 10X and 60X magnification. (B) To visualize DNA (green) in XDM/Coll I scaffolds, the scaffolds were stained with SYBR green dye and imaged using confocal laser scanning microscopy. A higher magnification picture of the area marked by a white dashed line box is shown. (C) To visualize collagen type I (pink) in XDM/Coll I scaffolds, the scaffolds were stained with Direct red 80 dye and observed in a phase-contrast light microscope. A higher magnification picture of the area marked by a white dashed line box is shown.

We further investigated if the scaffolds formed resulted from the combination of both DNA and Coll I. Therefore, we treated the scaffolds with SYBR green and picrosirius red to selectively stain DNA and Coll I, respectively. We observed that 20% and 50% XDM/Coll I scaffolds were extensively stained with SYBR green and picrosirius red, indicating that the scaffolds were made up of both DNA and collagen **(Fig. 2)**. Further, SYBR green staining demonstrated that the 20% XDM/Coll I scaffolds were made up of thin, extensive fibrous network, whereas 50 % XDM/Coll I scaffolds were made up of thicker fibrous network. Hence, the thickness of the individual fibrils can be regulated by modulating the mass fractions of XDM and Coll I. The flexibility to modulate the thickness of fibrils in a fibrous network can be an excellent strategy for developing next-generation highly tunable extracellular matrices (ECM) with thicknesses similar to that of native tissue, especially considering tissue with fibrous architecture^20–23^. For example, a fibrous network similar to that of nerve tissue can be developed for nerve tissue engineering applications. Furthermore, the produced scaffolds have the potential to serve as a coating material for various scaffolds, especially implants with poor cellular adhesion, and can be tailored to grow depending on the types of cells or tissues utilized^24,25^.

### 2.3. Characterization of XDM/Coll I Scaffolds

To further characterize the nature and morphology of scaffolds in greater detail, SEM and AFM were performed. Similar to the optical microscope observations, SEM analysis revealed the fibrous nature of XDM/Coll I scaffolds **(Fig. 3A)**. The surface of both 20% and 50% XDM/Coll I scaffolds displayed continuous intertwined fibrous network with 20% XDM/Coll I scaffolds with finer fibrous network in comparison to 50% XDM/Coll I scaffolds. By contrast, the surface of the collagen control (0% XDM/Coll I) and DNA control (100% XDM/Coll I) displayed continuous deposition of Coll I and X-DNA on the glass coverslip with morphology different from both 20% and 50% XDM/Coll I scaffolds. Further suggesting the development of a unique morphology from the interaction of X-DNA and Coll I.

**Fig. 3.**
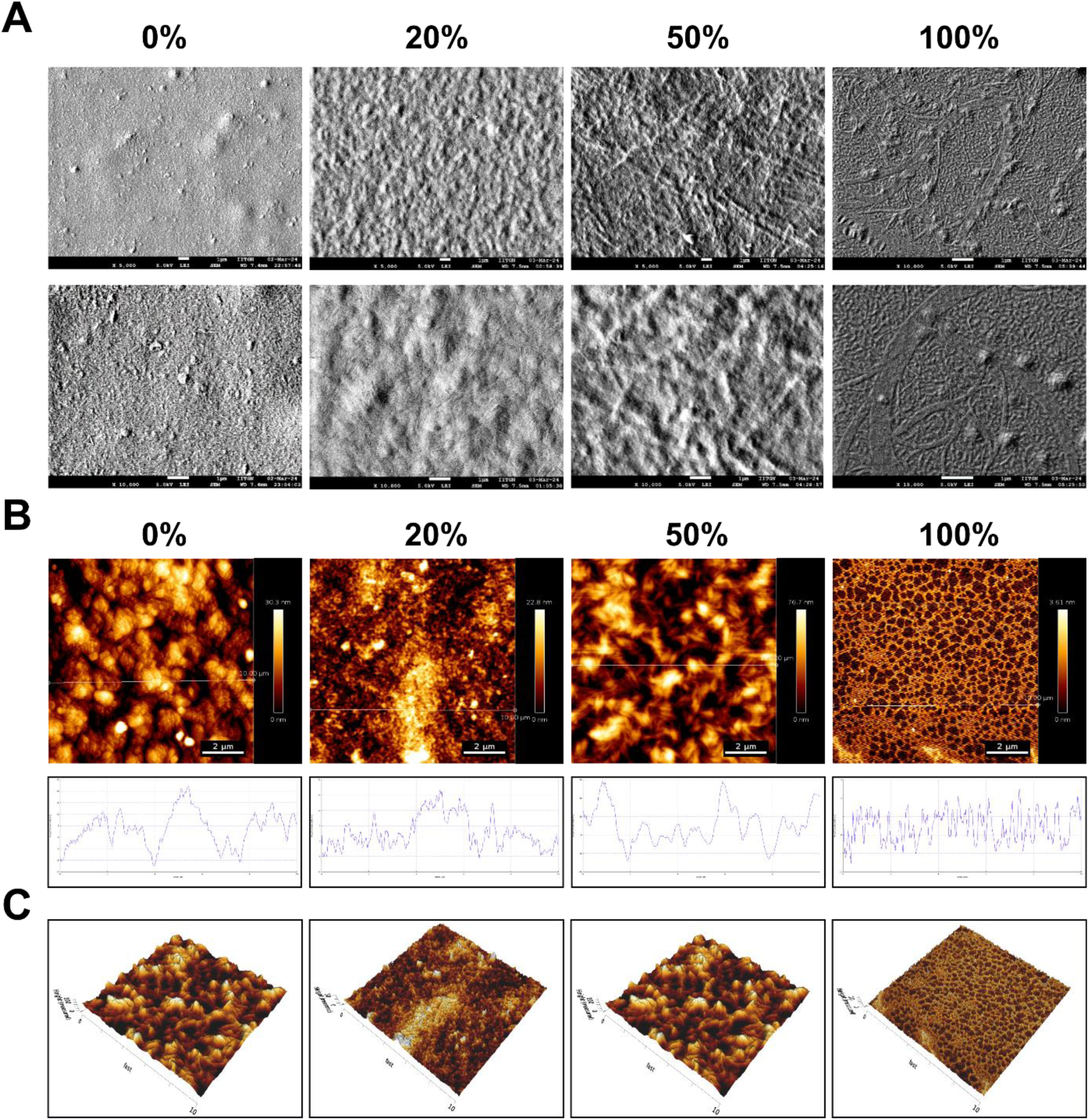
Characterization of XDM/Coll I scaffolds. (A) Field-emission scanning electron micrographs of control groups (0%, 100%) and XDM/Coll I scaffolds at increasing magnification. (B) AFM images of the samples with a plot displaying the height profile of the sample roughness in the region along the line are shown (line length: 10 µm). (C) The 3D profile of the AFM micrograph for different samples (height gradient bar for all samples is in the nm range).

To further visualize the morphology of the scaffolds at the nanoscale level with information about the height and 3D architecture, AFM imaging of the scaffolds was performed **(Fig. 3B)**. AFM characterization of the scaffolds revealed the highly intertwined fibrous network with 20% XDM/Coll I scaffolds having features with a maximum height of 22.8 nm, whereas 50% XDM/Coll I scaffolds had larger features with a maximum height of 76.9 nm, which is in line with the observation that 50% XDM/Coll I scaffolds had bigger fibrils in comparison to 20% XDM/Coll I scaffolds. Further, we observed that the fibrils formed in 50% XDM/Coll I scaffolds were made up of smaller fibrils intertwining together to form larger fibrils. We believe that as the amount of DNA increases, the fibrillogenesis of collagen around the DNA increases, resulting in the formation of larger fibrils in 50% XDM/Coll I scaffolds compared to 20% XDM/Coll I scaffolds. The 3D maps of the scaffolds further revealed the surface morphology and distribution of fibrils on the scaffolds **(Fig. 3C)**. Overall, the scaffolds formed had a fibrous architecture, different from X-DNA and Coll I alone. The fibrous architecture depended on the mass fraction of X-DNA and Coll I, with more prominent features observed as the mass fraction increased.

### 2.4. In Vitro Cell Culture on XDM/Coll I Scaffolds

To understand the physiological and biological effects of scaffolds on cells, SUM159 triple-negative breast cancer cells were cultured on scaffolds. We utilized the SUM159 triple-negative breast cancer cell line because of its high proliferation rate and stability, allowing for rapid expansion in culture with similar phenotypes over multiple passages, ensuring consistency and reliability in experiments^26^. Control experiments were performed by culturing the cells directly onto the glass coverslip (CS), collagen control, and DNA control. To visualize the effect of scaffolds on cell morphology and cytoskeletal arrangement, the cells were stained with phalloidin to stain the filamentous actin (F-actin). We observed that cells grown on a glass coverslip and DNA control demonstrated random directional growth, with cells preferably growing in huge clumps **(Fig. 4A)**. In contrast, cells grown on DNA/Collagen scaffolds preferentially grew either individually or in small clumps. Cells grown in collagen control demonstrated an anisotropic pattern growth, which can be attributed to cells growing along the collagen fibrils. However, 50% of XDM/Coll I scaffolds demonstrated an aligned cellular growth, with cells preferentially growing along the large fibrils on scaffolds. In contrast, the cells on 20% XDM/Coll I scaffolds didn’t demonstrate such extensive growth along the fibrils, indicating that the size of the fibrils in the fibrous network of scaffolds affects cellular growth patterns, with cells preferentially growing on large fibrils. We further quantified the mean fluorescent intensity of F-actin and observed that the cells on glass coverslips demonstrated the highest mean fluorescent intensity, whereas 20% and 50% XDM/Coll I scaffolds demonstrated significantly lower F-actin mean fluorescent intensity in comparison to glass coverslips **(Fig. 4C)**. We also observed that the cells on glass coverslips and DNA control developed significantly higher proportion of stress fibers which were significantly reduced in case of 20% and 50% XDM/Coll I scaffolds. We speculated that the decrease in F-actin mean fluorescent intensity on scaffolds with a decreased amount of F-actin stress fibers could be attributed to the scaffolds acting as a soft substrate for cells. Previous studies have demonstrated the role of matrix stiffness in modulating cellular attachment and cytoskeleton network with a stiffer matrix, resulting in higher F-actin organization than the softer matrix^27,28^. Therefore, glass coverslips, which provide a stiff matrix to cells, demonstrated higher F-actin mean fluorescent intensity compared to other groups. Overall, 20% and 50% XDM/Coll I scaffolds supported cellular growth, with 50% XDM/Coll I scaffolds resulting in aligned cellular growth preferentially along the large fibrils on the scaffolds. Also, the scaffolds acted as a soft substrate for cellular growth, resulting in lower F-actin organization.

**Fig. 4.**
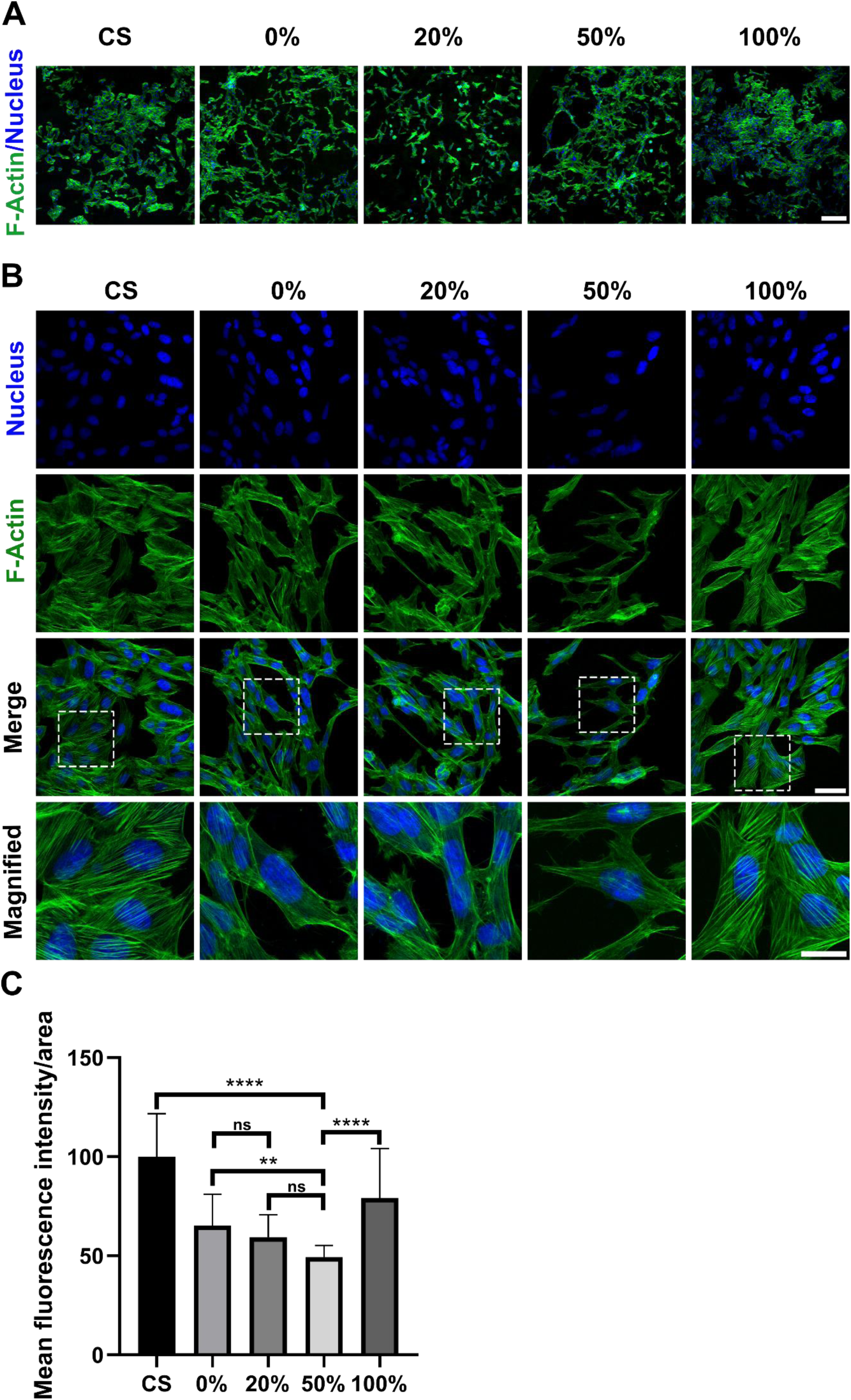
*In vitro* cell culture on XDM/Coll I scaffolds. Representative confocal microscope images for visualization of the growth pattern and cellular shape of SUM159 cells cultured on control groups and XDM/Coll I scaffolds by phalloidin staining (green) of F-actin at (A) lower magnification (10X objective: scale bar – 250 µm) and (B) higher magnification (63X oil objective: scale bar – 50 µm). Nuclei (blue) are stained with DAPI. For F-actin stress fibers visualization, the zoomed-in section includes a magnified image of the area marked with a white box in the merged groups (Scale bar - 25 µm) (C) Quantitative analysis of the mean fluorescent intensity per unit area of F-actin using image analysis. For quantification, a total of n=30 cells were randomly selected from different regions of the sample. Data presented as mean ± SD. * indicate statistical significance within respective groups. **, **** indicate p<0.01, and p<0.0001 respectively, and ns indicates no significant difference.

### 2.5. Effect of XDM/Coll I Scaffolds on Cellular Attachment

XDM/Coll I scaffolds acting as a soft substrate for cell growth warrant further investigation, especially considering that the actin cytoskeleton dynamics are directly linked to ECM via adhesion molecules and are involved in transmitting various biochemical signals^29,30^. We speculated that the lower F-actin organization observed in cells grown on scaffolds may be due to lower cellular adhesion of cells to soft substrate scaffolds. Therefore, we investigated the levels of Integrin-β1 and vinculin, the main focal proteins involved in cellular adhesion^31–33^. As expected, immunocytochemistry revealed a significantly higher expression of both integrin-β1 and vinculin in cells grown on glass coverslips and DNA control, indicating that the cells grown on these substrates are experiencing higher stiffness, with DNA control having lower stiffness in comparison to glass coverslips **(Fig. 5A,5B)**. In contrast, scaffolds of collagen control, 20% XDM/Coll I and 50% XDM/Coll I, act as soft substrates as indicated by significantly lower expression of integrin-β1 and vinculin in comparison to a glass coverslip and DNA control. Also, well-developed adhesion plaques of vinculin were observed on glass coverslip and DNA control, which were poorly observed on 20% and 50% XDM/Coll I scaffolds **(Fig. 5B)**. However, among all the groups, the 50% XDM/Coll I scaffolds demonstrated the lowest expression of integrin-β1 and vinculin, which agrees with their lowest F-actin expression among all groups, indicating that 50% XDM/Coll I scaffolds may have the lowest stiffness among all the tested groups.

**Fig. 5.**
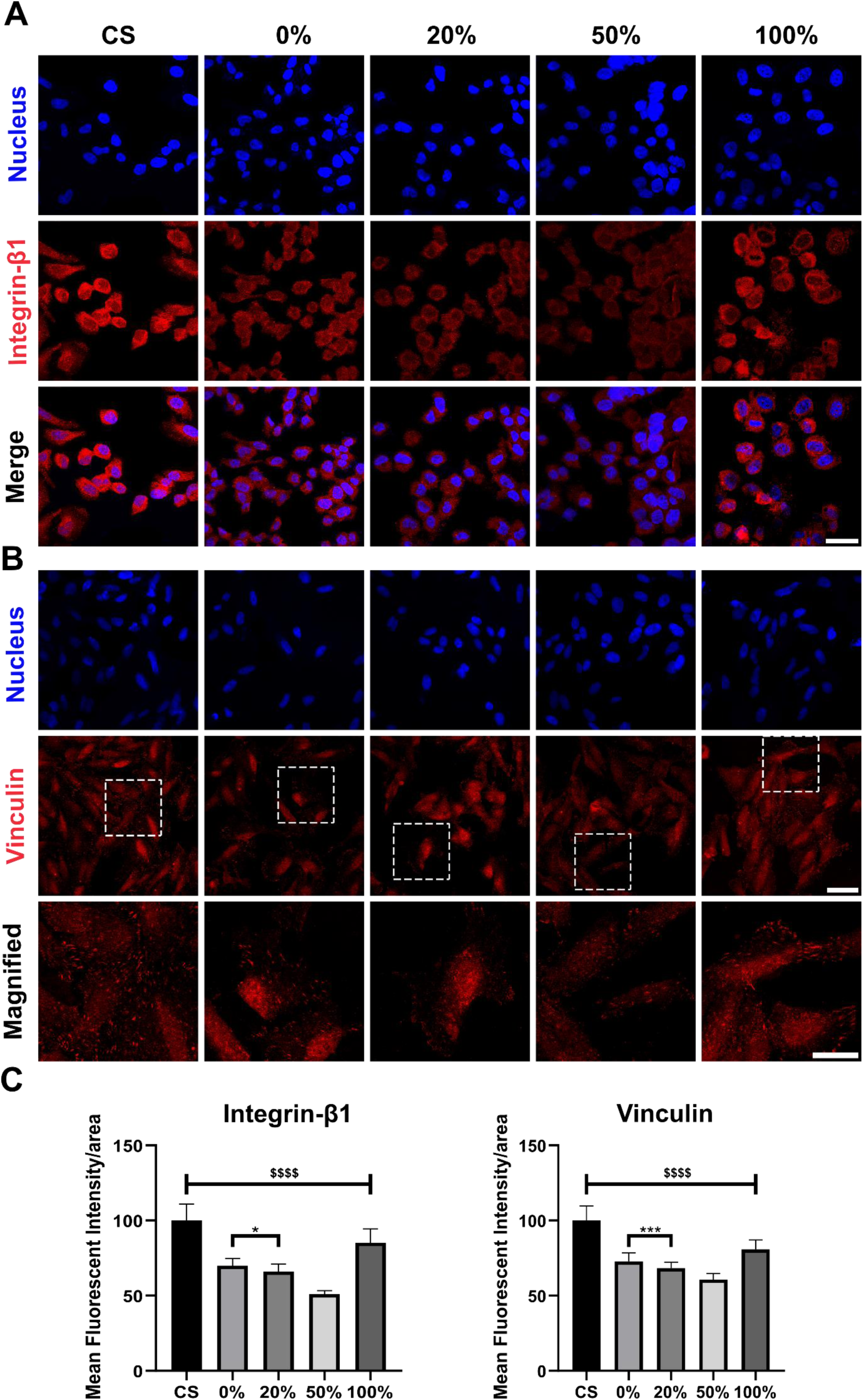
Effect of XDM/Coll I scaffolds on cellular attachment markers. Immunofluorescence staining of SUM159 cells cultured on control groups and XDM/Coll I scaffolds for (A) Integrin-β1 (red) and (B) vinculin (red) and nuclei (blue) (scale bar – 50 µm). For visualization of vinculin focal adhesion plaque, the zoomed-in section includes a magnified image of the area marked with a white box in the vinculin groups (Scale bar - 25 µm) and (C) Quantitative analysis of the mean fluorescent intensity per unit area of integrin-β1 and vinculin using image analysis. For quantification, a total of n=30 cells were randomly selected from different regions of the sample. Data presented as mean ± SD. *, $ indicate statistical significance within respective groups and 50% XDM/Coll I scaffolds, respectively. *, ***, $$$$ indicate p<0.05, p<0.001, and p<0.0001, respectively.

### 2.6. Effect of XDM/Coll I Scaffolds on cellular uptake of endocytic cargo

Internalization of extracellular materials through endocytosis is an essential process of the cell and is involved in many biological processes, such as cell survival, proliferation, differentiation, signal transduction, etc^34–36^. Also, enhanced cellular uptake of therapeutics, especially nanoparticles, has been a primary focus of many drug delivery strategies^37^. However, significant research efforts have been directed toward enhancing cellular uptake by modifying the characteristics of cargo, including their shape, dimensions, and surface charge, and limited utilization of cells and their ECM for enhancing cellular uptake has been studied^38–40^. In a recent study by Cassani et al., the authors demonstrated that cells grown on a softer substrate coated with collagen had an enhanced nanoparticle uptake compared to cells grown on a stiffer polystyrene substrate^41^. Similarly. the study by Alharbi et al. demonstrated an inverse correlation between endocytic processes and focal adhesion dynamics^42^. Hence, we next investigated the effect of soft 20% and 50% XDM/Coll I scaffolds on cellular uptake. For our investigation, we utilized Transferrin (Tf), a classical ligand that gets readily endocytosed by a clathrin-mediated pathway^43^. As endocytosed vesicles mature from early endosomes to late endosomes over a while, we investigated cellular uptake of transferrin at three different intervals: 15 min, 45 min, and 120 min. We compared the percentage uptake of transferrin in all conditions relative to glass coverslip with cellular uptake in glass coverslip normalized to 100%. We observed increased cellular uptake of transferrin in all other experimental groups to glass coverslips, which follows the understanding that the cellular uptake on stiffer matrices is lower in comparison to softer matrixes **(Fig 6A)**. Both 20% and 50% XDM/Coll I scaffolds resulted in significantly higher cellular uptake at all time intervals in comparison to other control groups, with 50% XDM/Coll I scaffolds resulting in cellular uptake 2 times and 3 times higher than glass coverslip at 15 min and 120 min, respectively **(Fig 6B)**. These results demonstrate the potential of XDM/Coll I scaffolds to be utilized as a material for enhancing the cellular uptake of cargo such as drugs, proteins, and small molecules without modifying the cargo itself. Moreover, utilizing XDM/Coll I scaffolds with cargo modified for enhanced cellular uptake can result in a multi-fold increase in the cellular uptake of cargo. Overall, 20% and 50% XDM/Coll I scaffolds demonstrated significantly higher cellular uptake of transferrin in comparison to the control groups. This can be attributed to these scaffolds serving as softer substrates, which are known to facilitate increased cellular uptake compared to stiffer matrices.

**Fig. 6.**
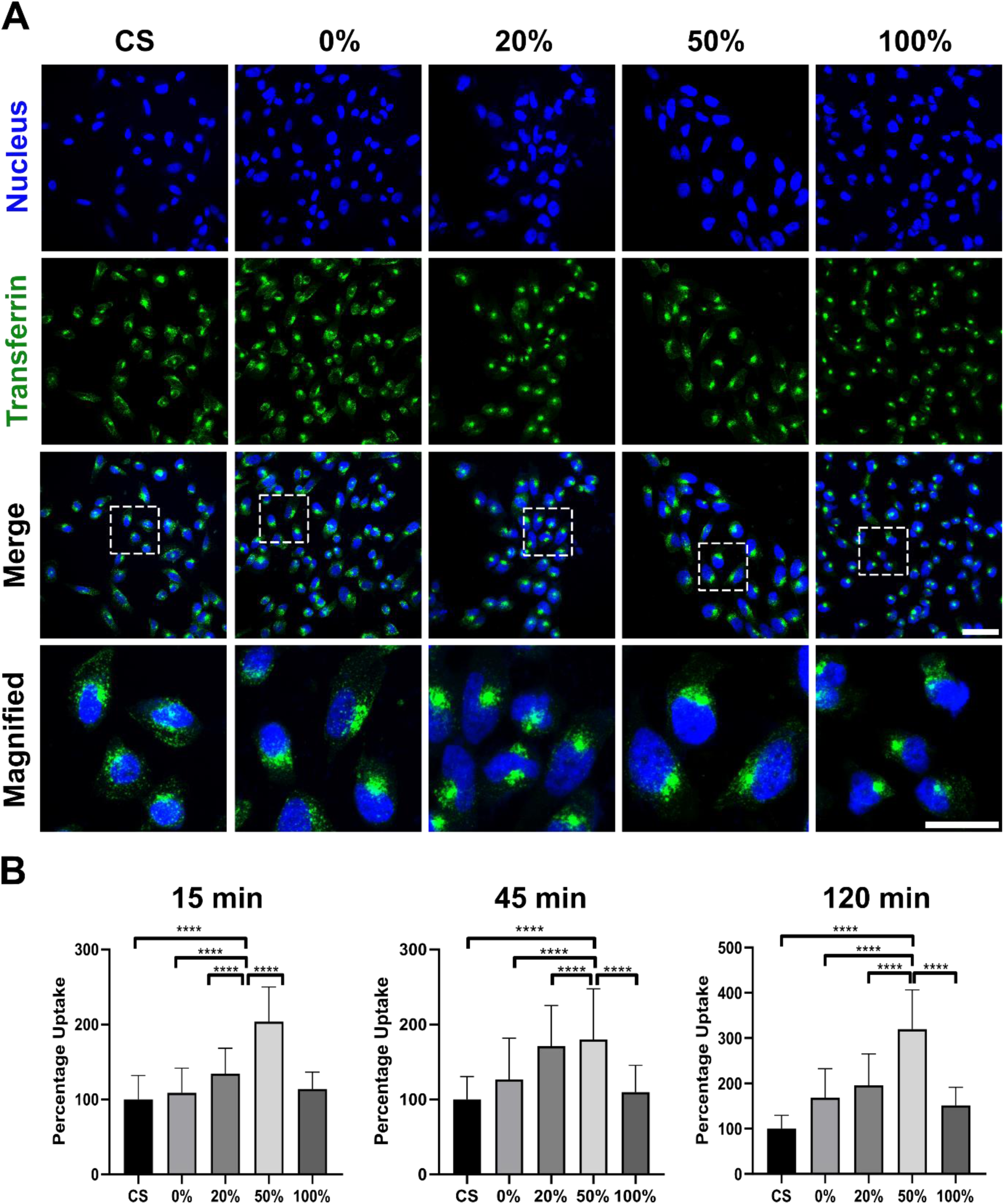
Effect of XDM/Coll I scaffolds on cellular uptake. (A) Representative confocal microscope images of SUM159 cells treated with FITC-conjugated transferrin (green) for 120 min on XDM/Coll I scaffolds (scale bar – 50 µm). A zoomed-in section includes a magnified image of the area marked with a white box in the merged groups (scale bar - 25 µm) (B) Quantitative analysis of the fluorescent intensity for percentage uptake of transferrin in SUM159 cells at different time points (15 min, 45 min, and 120 min) using ImageJ image analysis. For quantification, a total of n=30 cells were randomly selected from different regions of the sample. Data presented as mean ± SD. * indicate statistical significance within respective groups. **** indicate p<0.0001.

### 2.7. Effect of XDM/Coll I Scaffolds on Cellular Differentiation of SH-SY5Y Neuroblastoma Cells

A major focus of stem cell tissue engineering is to develop bioactive scaffolds that can not only support cellular growth but also provide necessary cues for the differentiation of stem cells into desired lineages^44,45^. *In situ,* autologous transplantation of stem cells at the site of injury has been utilized in many clinical trials for the treatment of different diseases^46,47^. These stem cells are culture-expanded and differentiated *in vitro* to the required cell density for successful implantation. However, the differentiation of stem cells into specific lineages with high efficiency utilizing minimum growth media components in a short period is still a critical aim of tissue engineering^48^. These limitations are especially evident while differentiating precursor neuronal cells into specific subtypes, as they require not only multiple neurotrophic growth factors over a long period but also an appropriate ECM environment with specific stiffness^49,50^.

We hypothesized that the soft matrix of XDM/Coll I scaffolds, along with its fibrillar architecture, is similar to the native ECM of nerve tissue and, thus, may provide an appropriate environment for neuronal cell viability and differentiation. To test our hypothesis, we cultured the SH-SY5Y human neuroblastoma cell line, which can differentiate into mature neurons on XDM/Coll I scaffolds. SH-SY5Y cells were cultured on glass coverslips, collagen control, DNA control, and 20% and 50% XDM/Coll I scaffolds with differentiation media. To observe the growth and morphology of cells, optical microscope images were taken every day till day 4, after which cells were fixed **(Fig 7A)**. From optical microscopy images, we observed poor growth of SH-SY5Y cells on a glass coverslip, which is expected as proper culturing of SH-SY5Y cells *in vitro* on tissue culture plates (TCP) requires ECM coating of TCP. Similarly, no change in the morphology of SH-SY5Y cells was observed for cells grown on DNA control. This can be due to SH-SY5Y cells experiencing a stiffer matrix, which is not similar to native nerve tissue and thus lacks the functionality to provide appropriate signals for cell growth and differentiation. In contrast to a glass coverslip and DNA control, cells grown on collagen control and 20% and 50% XDM/Coll I scaffolds demonstrated significant cellular attachment with spread morphology after 48 hrs only, indicating that the matrix of collagen control and scaffolds were favorable for cellular growth and viability (**Fig. S2A)**. However, from optical microscopy observation, we observed that cells grown on collagen control demonstrated a higher proliferation rate in comparison to both 20% and 50% XDM/Coll I scaffolds and were highly confluent at day 6 compared to 20% and 50% XDM/Coll I scaffolds. This observation indicates that the SH-SY5Y cells on collagen scaffolds may be undifferentiated as undifferentiated SH-SY5Y cells rapidly proliferate, while differentiated cells have a reduced proliferation rate^49^. Also, undifferentiated SH-SY5Y cells are non-polarized, while differentiated SH-SY5Y cells extend long and have branched processes/neurites. Therefore, we next observed the cytoskeleton of SH-SY5Y cells by fixing them on day 6 and staining them with phalloidin to mark F-actin. We observed an extensive change in the morphology of cells grown on 20% and 50% XDM/Coll I scaffolds, which was not observed in the control groups **(Fig 7B)**. The SH-SY5Y cells on 20% and 50% XDM/Coll I scaffolds acquired an elongated shape with visible polarity, a characteristic of differentiated SH-SY5Y cells. Additionally, extensive branched processes/neurites were observed on cells grown on 20% and 50% XDM/Coll I scaffolds, suggesting that the cells on 20% and 50% XDM/Coll I scaffolds have differentiated into neurons. To confirm the differentiation status of SH-SY5Y cells, we immunostained SH-SY5Y cells with primary antibody against Microtubule-associated protein-2 (MAP2), a protein involved in the stability and regulation of microtubule networks in axons and dendrites^51^. We observed several neurites extending from the SH-SY5Y cells grown 20% and 50% XDM/Coll I scaffolds, confirming that the cells have differentiated **(Fig. 7B)**. Also, very few neurites were observed in control groups compared to 20% and 50% XDM/Coll I scaffolds, indicating that the cellular differentiation of SH-SY5Y cells was a characteristic feature of DNA/collagen scaffolds. Also, the differentiation of SH-SY5Y cells on 20% and 50% XDM/Coll I scaffolds was achieved in 6 days without multiple passaging, indicative of the higher efficiency of XDM/Coll I scaffolds for SH-SY5Y differentiation. Quantification of neurite length demonstrated that the neurite length in 20% and 50% XDM/Coll I scaffolds were significantly longer than collagen control **(Fig. 7C)**. Among 20% and 50% XDM/Coll I scaffolds, 20% XDM/Coll I scaffolds resulted in higher neurite length, but the difference was not significant. However, we observed that the cells on 20% XDM/Coll I scaffolds formed more neurite junctions, whereas cells on 50% XDM/Coll I scaffolds preferentially grew along the large fibrils and formed an extensive, highly connected neuronal cell network along the fibrils **(Fig. S2B).** Overall, 20% and 50% XDM/Coll I scaffolds resulted in extensive differentiation of SH-SY5Y cells and thus can be the material of choice for achieving enhanced neuronal differentiation.

**Fig. 7.**
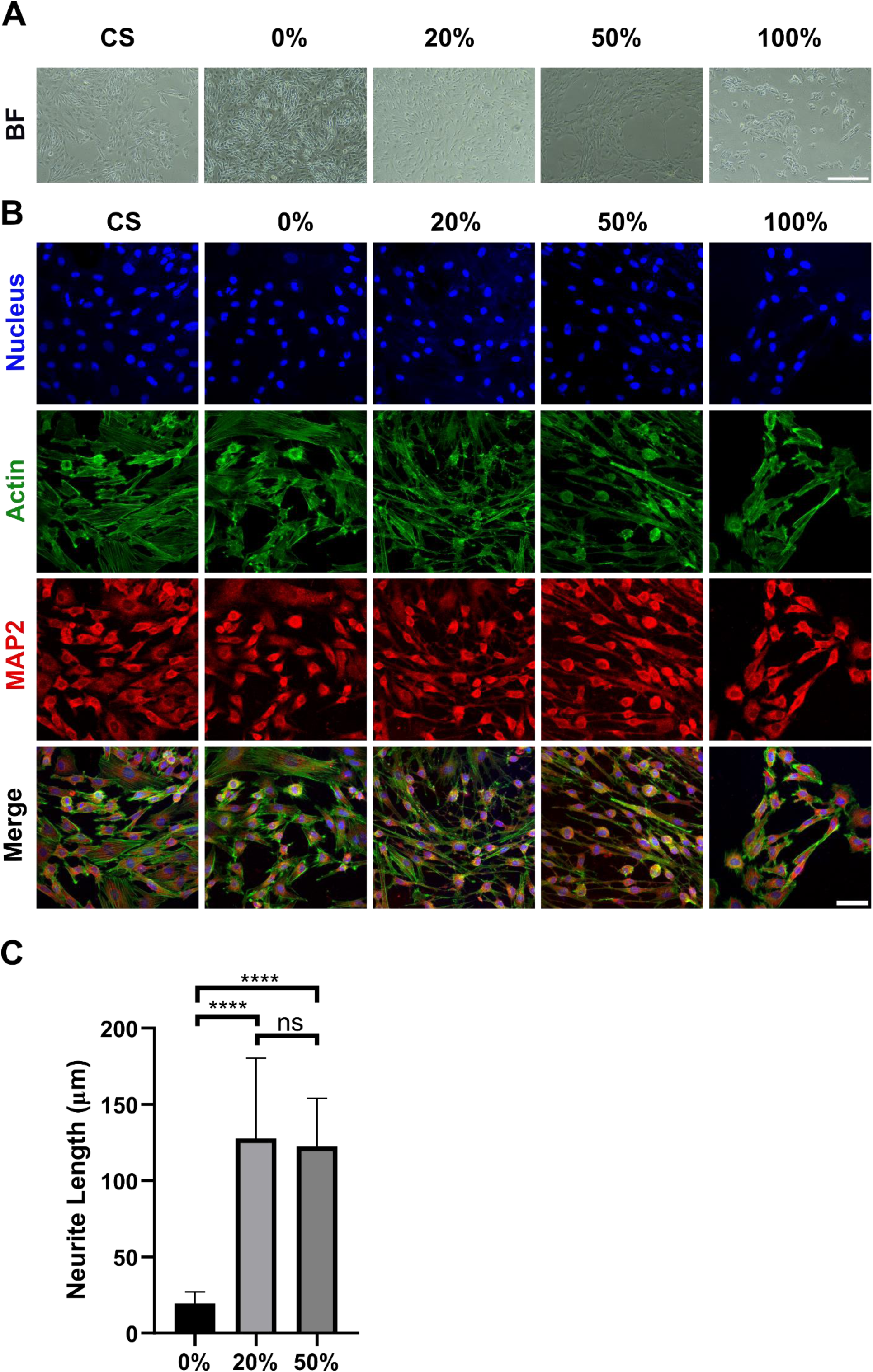
Differentiation of precursor SH-SY5Y cells using XDM/Coll I scaffolds. (A) Representative bright field images of SH-SY5Y cells cultured on control groups and XDM/Coll I scaffolds on day 6 (scale bar-250 µm). (B) Visualization of the cytoskeleton and differentiation status of 6-day cultured SH-SY5Y cells using phalloidin staining (green) to stain F-actin and MAP2 immunostaining (red) to mark differentiated neurons (scale bar – 50 µm). (C) Quantitative analysis of the neurite length using ImageJ image analysis (NeuronJ). For quantification, a total of n=60 cells were randomly selected from different regions of the sample. Data presented as mean ± SD. * indicate statistical significance within respective groups. **** indicates p<0.0001, and ns indicates no significant statistical difference.

## 3. Conclusions

We report for the first time the interaction of self-assembled DNA macrostructure with collagen type I to form bioactive scaffolds. We investigated the effect of different mass fractions of XDM and collagen type I on scaffolds formation and observed that 20% and 50 % mass fractions resulted in bioactive scaffolds formation. The formed scaffolds were made up of dense intertwined fibrous networks, with 20% XDM/Coll I scaffolds containing thinner fibrils compared to 50% XDM/Coll I scaffolds, indicating that as the amount of DNA increases, the thickness of fibrils on scaffolds increases. *In vitro* cell culture experiment on XDM/Coll I scaffolds demonstrated that the scaffolds support cellular growth, with aligned cellular growth along the scaffolds fibrils observed on 50% XDM/Coll I scaffolds. We observed that XDM/Coll I scaffolds acted as a softer matrix for cell growth and had low F-actin organization with decreased expression of cell adhesion markers. Also, due to XDM/Coll I scaffolds serving as softer substrates, they demonstrated significantly higher cellular uptake of transferrin in comparison to the control groups. Lastly, XDM/Coll I scaffolds supported neuronal cell growth and resulted in the differentiation of SH-SY5Y neuroblastoma cells within 4 days. The SH-SY5Y cells on XDM/Coll I scaffolds demonstrated significant cell polarity with long neural projection, which was not observed in control groups. Overall, we report the formation of bioactive scaffolds made from the interaction of self-assembled DNA macrostructure and collagen type I, which can be utilized for diverse biomedical applications, including *in vitro* cell culture for specialized cell growth, drug delivery for enhanced cellular uptake, stem cell tissue engineering for differentiating stem cells into specific lineages and as a coating material for various scaffolds to improve cellular growth and differentiation.

## 4. Materials and Methods

### 4.1. Reagents

All DNA primers were purchased from Sigma-Aldrich, USA, with 0.2 µmole scale synthesis and desalting purification. Nuclease-free water (Sisco Research Laboratory), magnesium chloride (SRL, India), 6X gel loading dye (HiMedia), collagen type I rat tail (Gibco), Direct Red 80 (Sigma-Aldrich, USA), Dulbecco’s modified Eagle’s medium (DMEM: Gibco), Ham’s F-12 Nutrient Mixture (Gibco), Fetal bovine serum (FBS: Gibco), Transferrin (Tf: Santa Cruz Biotechnology), Integrin-Beta (Invitrogen: 4706S), Vinculin (Invitrogen: 44-1074G), Bovine serum albumin (BSA: Sigma-Aldrich, USA), Glycine (HiMedia). Alexa Flour 488 phalloidin (Thermo Fisher Scientific), B-27 (Gibco), Glutamax (Gibco), and Retinoic acid (TCI chemicals)

### 4.2. Preparation of the Self-assembled DNA Macrostructure

The X-DNA macrostructure (XDM) was assembled using four ssDNA primers (**Table S1**: ssDNA sequence) following the previously published protocol^52^. Briefly, four oligonucleotides, X1, X2, X3, and X4, were mixed in an equimolar ratio in nuclease-free water with 2 mM MgCl_2_. The macrostructure was assembled by heating the primers at 95 °C for 30 min and then cooling them to 5 °C with a step decrease of 5 °C every 15 min using a PCR instrument (BioRad, USA). The resulting macrostructure was then stored at 4 °C until required. Different molar concentrations of the primers were utilized depending on the mass fraction of collagen to DNA, with a total mass of 102.66 µg. An Electrophoretic Mobility Shift Assay (EMSA) was performed to confirm the formation of XDM. Briefly, a 10% native polyacrylamide gel was prepared, and the wells were loaded with 2 µl of DNA samples, along with 2 µl 6X gel loading dye. The gel was run at a constant voltage of 90 V for 60 min, followed by ethidium bromide staining for 20 min on a shaker. The gel was then imaged using a Gel Documentation system (BioRad ChemiDocTM MP Imaging System).

### 4.3. Formation of DNA-Collagen Scaffolds

XDM solution was mixed with collagen type I (Coll I) to obtain different mass fractions (XDM / XDM + Coll I) of 0%, 20%, 50%, and 100%, with a total mass of 102.66 µg and final volume of 200 µL **(Table S2).** The mass fraction of 0% and 100% was taken as the collagen control and XDM control, respectively, in all experiments. After mixing, the solution was rigorously vortexed for 2 min. The resulting solution was subsequently transferred to a 10 mm glass coverslip in a 24-well tissue culture plate and vacuum dried for 48 h at room temperature (25 °C) to form the XDM/Coll I scaffolds.

### 4.4. Atomic Force Microscopy

The XDM/Coll I scaffolds on glass coverslips were glued to a glass slide and analyzed directly. The images were collected using an Atomic Force Microscope (Bruker) in air-tapping mode using a cantilever tip (Force Mod, 8 nm Tip radius, 80 KHz).

### 4.5. Scanning Electron Microscopy

The XDM/Coll I scaffolds on glass coverslips were taped to copper stubs using double-sided carbon tape. The scaffolds were sputter-coated with platinum for 60 s (10 mA, 5 mTorr). Images were collected using field emission SEM (JEOL JSM-7600F, Tokyo, Japan) at an acceleration voltage of 5 keV.

### 4.6. Picrosirius Red Staining

The XDM/Coll I scaffolds were hydrated with deionized water for 3 min and then stained with picrosirius red stain (1 mg/mL of Direct red 80 in saturated picric acid solution) for 60 min at room temperature (25 °C). Further, the scaffolds were washed with acidified water (0.5% acetic acid solution) 3 times. The scaffolds were dehydrated with absolute ethanol for 5 min, and stained scaffolds were imaged by phase contrast microscopy (Nikon Eclipse Ts2).

### 4.7. Cell Culture

SUM159 triple-negative breast cancer cell lines were used for cell growth and proliferation studies. SUM159 cells were suspended in Ham’s F-12 Nutrient Mixture (F-12) media with 5% FBS and 1% antimycotic-antibiotic (penicillin-streptomycin) solution. The cells were grown at 37 °C in a cell culture incubator at 5% CO_2_. For the growth and proliferation study, cells were seeded on XDM/Coll I scaffolds with a seeding density of 12.7 × 10^3^ cells/cm^2^ for 3 days. After 3 days, the cells were fixed with 4% paraformaldehyde and stained with Alexa Flour 488 phalloidin. For nucleus staining, the coverslips with fixed scaffolds were mounted on glass slides using Mowiol containing 10µg/mL DAPI. Images were obtained using a 63X oil objective on a confocal laser microscope (Leica TCS SP8).

### 4.8. Assessment of Cellular Uptake Using Transferrin

To study the cellular uptake of transferrin, SUM159 cells were seeded on XDM/Coll I scaffolds with a seeding density of 12.7 × 10^3^ cells/cm^2^. Cells were maintained for 3 days till 70-80% confluency was reached. Cells were treated with 5 µg/mL Transferrin (Tf) and fixed using 4% paraformaldehyde at three different time points (15 min, 45 min, and 120 min). The glass coverslips containing fixed scaffolds were then mounted using Mowiol containing DAPI and imaged using a 63X oil objective on a confocal laser microscope.

### 4.9. Immunohistochemistry

Immunohistochemistry of cells grown on XDM/Coll I scaffolds was performed to visualize Integrin-Beta, vinculin, and MAP2 expression using antibodies against them as per manufacturer protocol. Briefly, SUM159 cells seeded on XDM/Coll I scaffolds at a seeding density of 12.7 × 10^3^ cells/cm^2^ were fixed on day 3 using 4% paraformaldehyde for 15 min and subsequently permeabilized using 0.1% Triton X- 100 for 10 min. Permeabilized cells were then blocked using blocking buffer (1% BSA, 300 mM Glycine, 0.1% Tween 20 in 1X PBS) for 60 min at room temperature. Next, the fixed samples were stained with primary antibodies against integrin-beta and vinculin for 2 h in a humidified chamber at room temperature (25 °C). The primary antibody was diluted at 1:250 dilution in antibody dilution buffer (1% BSA in PBST (1X PBS + 0.1% Tween 20)). The cells were then labeled with 4 µg/mL goat anti-rabbit (For Integrin-Beta and Vinculin: Invitrogen; A11011) or anti-mouse (For MAP2: Sigma Aldrich; SAB4700503) IgG secondary antibody for 2 h at room temperature in the dark. The samples were mounted using Mowiol containing DAPI and imaged using a 63X oil objective on a confocal laser microscope.

### 4.10. Differentiation of SH-SY5Y cells

Differentiation of SH-SH5Y cells was carried out following a previously published protocol^52^. Briefly, undifferentiated precursor SH-SH5Y cells were seeded on XDM/Coll I scaffolds with a seeding density of 12.7 × 10^3^ cells/cm^2^. The cells were allowed to attach and grow on scaffolds in basic growth media (DMEM/F-12) for 2 days. After 2 days, basic growth media was replaced with differentiation media (1% B-27, 20 mM KCl, 2 mM Glutamaxl, 1% Pen/Strep, and 10 µM Retinoic acid), and media was changed every 24 h till day 6. Fresh 10 mM retinoic acid was added during each media change. After day 6, cells were fixed and stained for further analysis.

### 4.11. Statistical Analysis

Each experiment was performed independently at least 3 times, and the results have been presented as the mean ± standard deviation for each. Representative data of the 3 repeats has been provided. Statistical analysis was conducted using GraphPad Prism 8 software, employing one-way analysis of variance (ANOVA), followed by Tukey’s post-hoc analysis to determine statistical significance. Statistical significance was considered at p<0.05.

## Acknowledgments

The authors sincerely thank all of the members of the D.B. group for critically reading the manuscript and for their valuable feedback. The authors recognize IIT Gandhinagar’s infrastructure and financial support for the conduct of this research. NS and AS acknowledge the financial support from the Ministry of Education, Government of India. AS thanks PMRF fellowship. The authors acknowledge the Central Instrumentation Facility at IIT Gandhinagar for assistance with Field emission-SEM microscopy, AFM microscopy, and Confocal microscopy. D.B. thanks SERB, and GoI for the Core research grant, IITGN for the start-up grant, and Gujcost-DST, GSBTM, and STARS-MoES for research grants. D.B. is a member of the Indian National Young Academy of Sciences (INYAS).

## Authors contributions

Conceptualization: NS and DB; Methodology: NS, AS, and DB; Formal analysis and investigation: NS; Validation: NS and DB; Writing - original draft preparation: NS; Writing - review and editing: NS and DB; Funding acquisition: DB; Resources: DB; Supervision: DB; Project administration: DB.

### Data availability

All data generated or analyzed during this study are included in this published article.

