## Supporting information for "Self-assembled DNA-collagen bioactive scaffolds promote cellular uptake and neuronal differentiation"

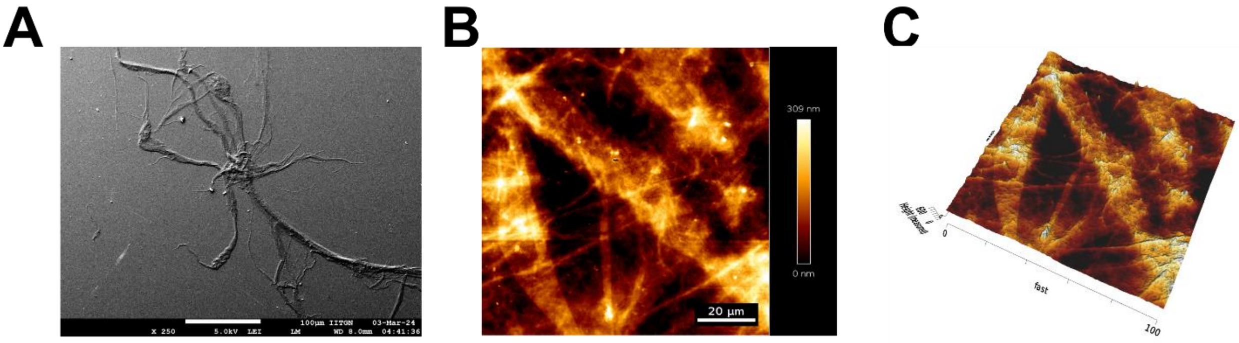


**Fig. S1** Characterisation of the fibers formed by the interaction of ssDNA primer (X1) and collagen type I. (A) Field-emission scanning electron micrographs of fibers. (B) AFM images of the fibres (scale bar: 20 µm). (C) The 3D profile of the fiber AFM micrograph (height gradient bar: 0 nm to 600 nm).

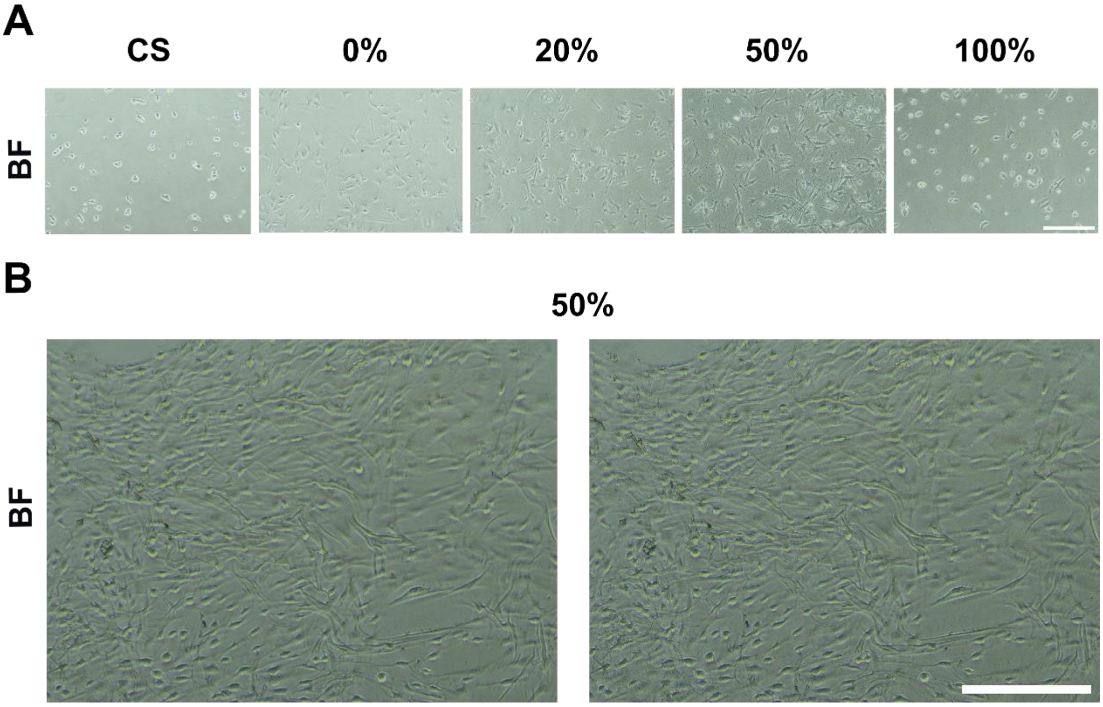


**Fig. S2** Representative bright field images of SH-SY5Y cells cultured on control groups and XDM/Coll I films after (A) 2 days post-seeding (scale bar- 250 µm). (B) An extensive, highly connected neuronal cell network of SH-SY5Y cells on 50% XDM/Coll I films at day 6 (scale bar- 250 µm).

**Table S1:** X-DNA macrostructure primer sequence

| ssDNA Primer | Sequence |
|---|---|
| X1 | 5'-CACGTGCGACCGATGAATAGCGGTCAGATCCGTACCTACTCG-3' |
| X2 | 5'-CACGTGCGAGTAGGTACGGATCTGCGTATTGCGAACGACTCG-3' |
| X3 | 5’-CACGTGCGAGTCGTTCGCAATACGGCTGTACGTATGGTCTCG-3' |
| X4 | 5'-CACGTGCGAGACCATACGTACAGCACCGCTATTCATCGGTCG-3' |

**Table S2:** Different Mass fraction ratio of XDM and Collagen type I

| Sample | DNA mass (µg) | Coll I mass (µg) | Total mass (µg) |
|---|---|---|---|
| 0% | 0 | 102.66 | 102.66 |
| 20% | 20.53 | 82.12 | 102.66 |
| 50% | 51.33 | 51.33 | 102.66 |
| 100% | 102.66 | 0 | 102.66 |
